# Computational Methods for Biofabrication in Tissue Engineering and Regenerative Medicine - a literature review

**DOI:** 10.1101/2023.03.03.530995

**Authors:** Roberta Bardini, Stefano Di Carlo

**Author notes:** Corresponding author (R. Bardini); (S.D. Carlo).

## Abstract

This literature review rigorously examines the growing scientific interest in computational methods for Tissue Engineering and Regenerative Medicine biofabrication, a leading-edge area in biomedical innovation, emphasizing the need for accurate, multi-stage, and multi-component biofabrication process models. The paper presents a comprehensive bibliometric and contextual analysis, followed by a literature review, to shed light on the vast potential of computational methods in this domain. It reveals that most existing methods focus on single biofabrication process stages and components, and there is a significant gap in approaches that utilize accurate models encompassing both biological and technological aspects. This analysis underscores the indispensable role of these methods in understanding and effectively manipulating complex biological systems and the necessity for developing computational methods that span multiple stages and components. The review concludes that such comprehensive computational methods are essential for developing innovative and efficient Tissue Engineering and Regenerative Medicine biofabrication solutions, driving forward advancements in this dynamic and evolving field.

**Graphical Abstract:** 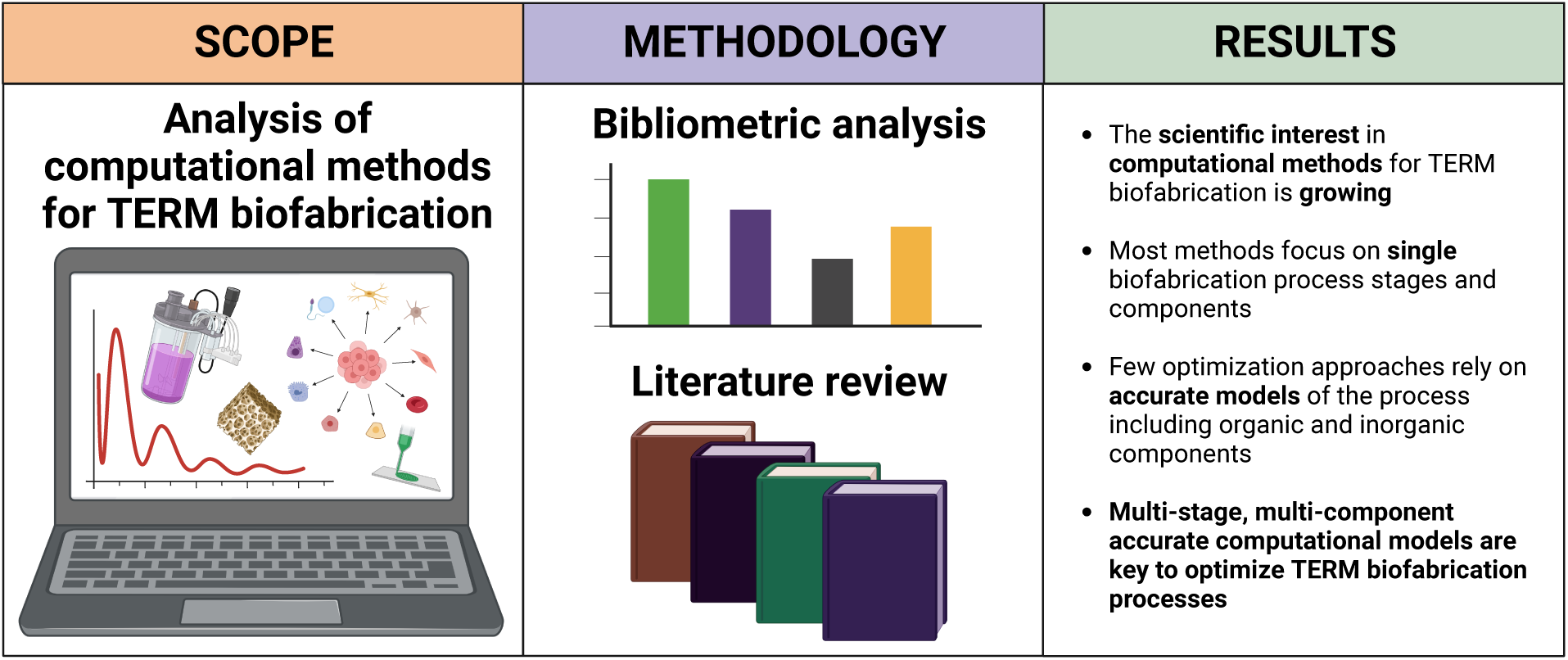

## 1. Introduction

Biofabrication is *"the automated generation of biologically functional products with the structural organization from living cells, bioactive molecules, biomaterials, cell aggregates such as micro-tissues, or hybrid cell-material constructs, through Bioprinting or Bioassembly and subsequent tissue maturation processes"* [1]. Tissue Engineering and Regenerative Medicine (TERM) is a challenging bio-fabrication application field bringing the promise to revolutionize the biomedical sector [2]. TERM applications require biofabrication products to be biomimetic, that is, to recapitulate the structural and functional features of their *in vivo* counterparts [3]. The degree of biomimesis in a product, based on the similarity of structural and functional features to physiological counterparts, determines its quality and clinical relevance. In biofabrication *"the process is the product"*, i.e., biofabrication processes and the resulting products are strictly intertwined, and so is their quality. Thus, ensuring product quality implies controlling process quality [4].

Quality control in TERM biofabrication requires analyzing the complex features determining the product quality, considering them as a function of defined critical process parameters [5]. Multi-technology biofabrication processes combine existing technologies to consider the product’s relevant scales and aspects, maximizing product quality [6]. Automation and digitalization optimize the execution of existing biofabrication processes or parts [7]. Finally, searching for improved or new process schemes moves from inefficient trial-and-error paradigms to intelligent design [8].

Computational approaches play a role in each of these tasks. They work to harmonize multi-technology schemes with automation, optimizing existing processes and supporting rational research design. This harmonization helps better organize the experimental activity to make process improvement more efficient [6]. Besides enabling existing technologies and experimental designs to work together, computational approaches have untapped potential to innovate biofabrication process designs. To truly impact biomimetic quality, intelligent design approaches must rely on accurate models of biofabrication [9, 10, 11].

This work analyses the application of intelligent automation principles (Figure 1.A) in modeling, design, and optimization (Figure 1.B), with a focus on approaches combining simulation and optimization (Figure 1.C) of TERM Bio-fabrication processes (Figure 1.D). The following sections delve into various aspects of the field. Initially, Section 2 presents a bibliometric analysis that sets the stage for understanding the current scientific landscape. This is followed by Section 3, which examines and evaluates the technological context relevant to computational methods in TERM bio-fabrication. Subsequently, Section 4 offers a comprehensive review of contemporary computational approaches in this domain, categorizing them according to the process stage they target. Finally, Section 5 synthesizes provided findings and discusses open challenges.

**Figure 1:**
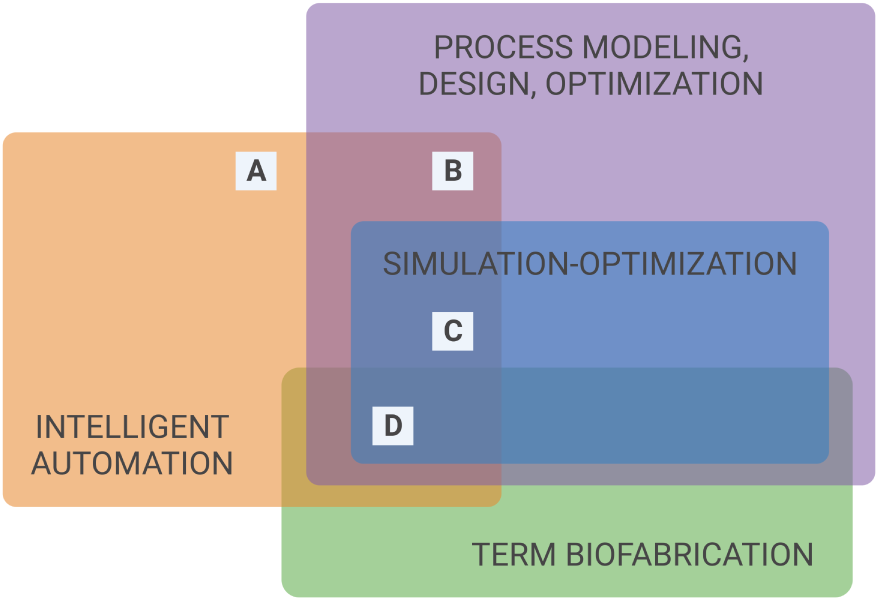
This work presents a review of the scientific literature on works that, following the principles of Intelligent Automation (A), leverage Process modeling, design and optimization (B), and in particular Simulation-optimization (C) techniques for TERM Biofabrication processes (D).

### Statement of significance

#### Issue

The central thrust of this review is to unveil the untapped potential of computational methods in enhancing the capabilities and outcomes of TERM biofabrication.

#### What is Already Known

Several computational approaches exist for different stages of the biofabrication process, and existing reviews analyze them thoroughly. Yet, existing analyses lack focus on the role of accurate models of the biofabrication process in computational approaches to TERM biofabrication.

#### What this Paper Adds

This work quantifies the relevance of the topic through the bibliometric and context analyses of the scientific domain of computational approaches to TERM biofabrication; subsequently, this work provides a review and analytical comparison of existing solutions organized by biofabrication process stage, including solutions based on accurate models of the biofabrication process.

## 2. The growing attention toward computational approaches: a bibliometric analysis

Attention toward computational methods is increasing in the biofabrication field. As illustrated in [12], several approaches exist to analyze a scientific landscape. This section enriches the study with a brief bibliometric analysis of the relevant scientific landscape. The analysis relies on PubMed^1^ queries that combine keywords related to Biofabrication, TERM, and computational approaches, plus synonyms and adjacent terms to search publications on the platform.

### 2.1. Performance analysis

This analysis starts with the claim that the scientific community’s interest in biofabrication and TERM is rapidly growing, supporting the relevance of the proposed work in analyzing state-of-the-art scientific production on the topic. Query 1.1 supports this claim by quantifing the interest in Biofabrication and Query 1.2 focuses the quantification on TERM Biofabrication. Results were analyzed considering the period between 2002 and 2021 and using the Total Publications (TP) count as a performance metric [12].

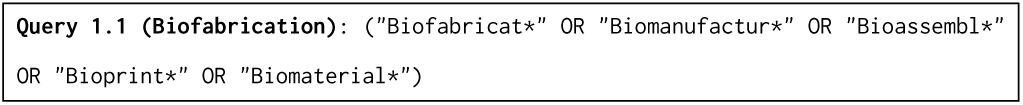

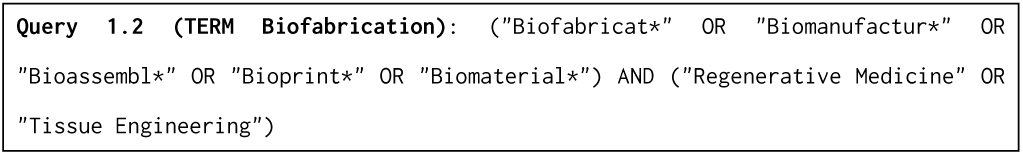

Query 1.1 retrieved 108,008 TP while Query 1.2 produced 28,549 TP, issued between 2002 and 2021. Figure 2 shows a steady increase in yearly TP during the last two decades, indicating a growing interest of the scientific community in the field. The term Biofabrication, in general, reported 1,221 TP in 2002, growing up to 12,921 TP in 2021 with a 958% increase. Of these publications, the portion on TERM Biofabrication grew from about 12% of the total (149 over 1,221 TP) in 2002 to about 27% in 2021 (3,516 over 12,921 TP).

**Figure 2:**
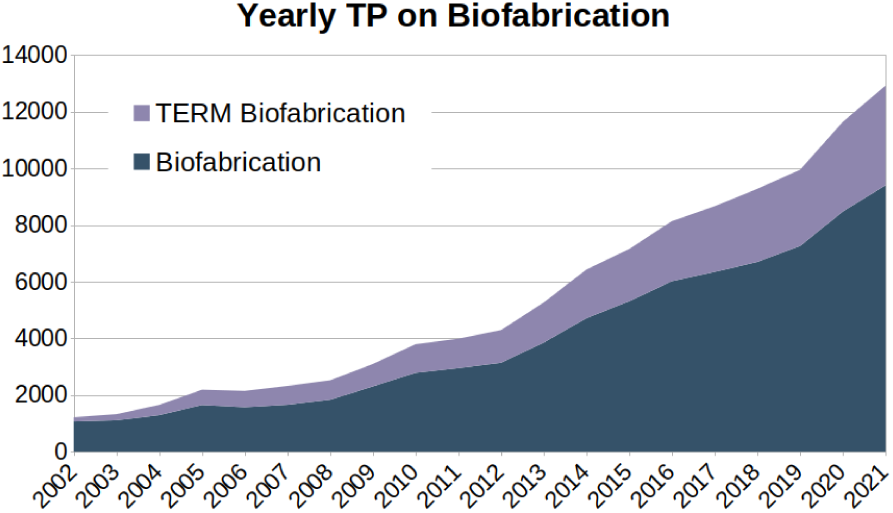
Total Publications per year on *Biofabrication*, with an highlight on the contributions on *TERM Biofabrication* (*Queries 1.1-1.2*).

Considering the 28,549 TP on TERM Biofabrication, the analysis considered publications on computational approaches to TERM Biofabrication (Query 2.1), and more specifically on computational approaches to TERM Biofabrication based on either modeling or optimization (Query 2.2), finally narrowing down the focus on computational methods to TERM Biofabrication based on combined modeling and optimization (Query 2.3).

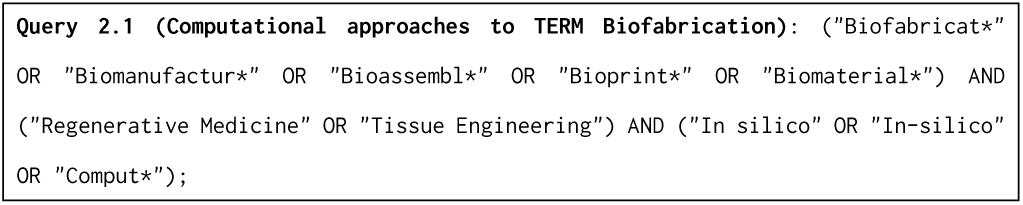

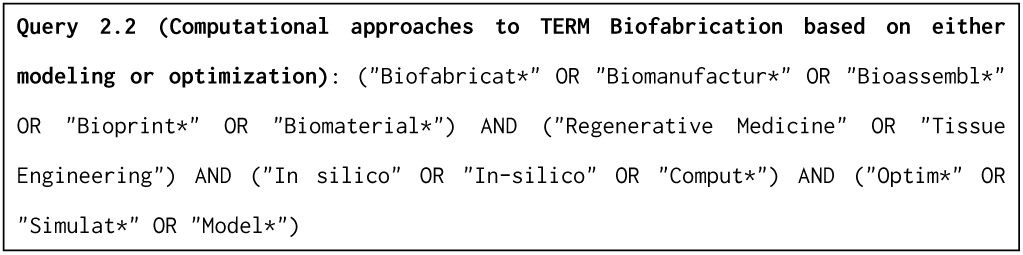

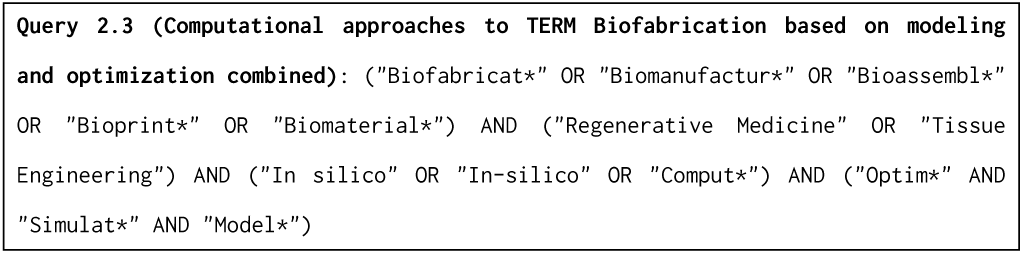

Starting from the result of Query 1.2, Queries 2.1-2.2 retrieved 1,642 TP for computational approaches to TERM Biofabrication (about 6%). Among these, 966 TP discuss computational approaches to TERM Biofabrication based on either modeling or optimization (about 60%). Figure 3 shows the TP per year for TERM Biofabrication, highlighting the presence of scientific activity on Computational approaches to TERM Biofabrication in time, which constitutes 5% (101 publications) of TP between 2002 and 2006, and 6% (785 publications) for the period between 2017 and 2021, as illustrated in Figure 4. Figure 5 shows that publications on computational approaches to TERM Biofabrication based on either modeling or optimization constitute 50% (50 over 100 publications) of TP on computational methods to TERM Biofabrication between 2002 and 2006 and the 57% (785 publications) for the period between 2017 and 2021.

Finally, Figure 6 illustrates the TP between 2002 and 2021 on computational approaches to TERM Biofabrication based on either modeling or optimization (Query 2.2, 966 TP), highlighting the contributions of computational methods to TERM Biofabrication based on combined modeling and optimization (Query 2.3).

**Figure 3:**
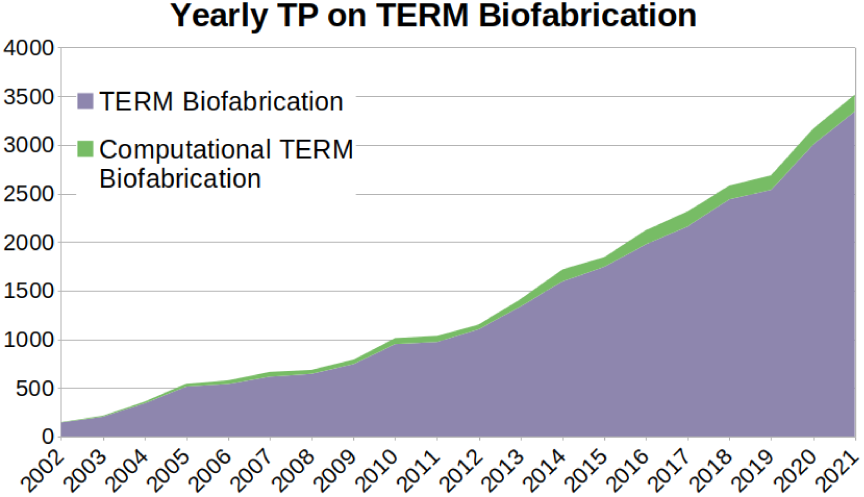
Total Publications per year on *TERM Biofabrication*, with an highlight on the contributions on *Computational approaches to TERM Biofabrication* (*Queries 1.2 and 2.1*).

**Figure 4:**
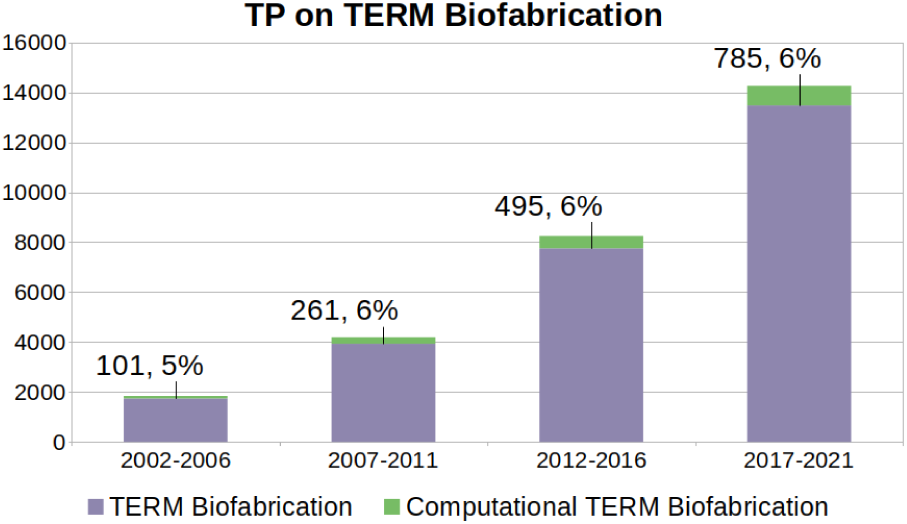
Total Publications on *TERM Biofabrication* aggregated over five-year periods, with a highlight on the contributions on *Computational approaches to TERM Biofabrication* (*Queries 1.2 and 2.1*), that constitute between 5-6% of TP across all periods from 2002 to 2021.

**Figure 5:**
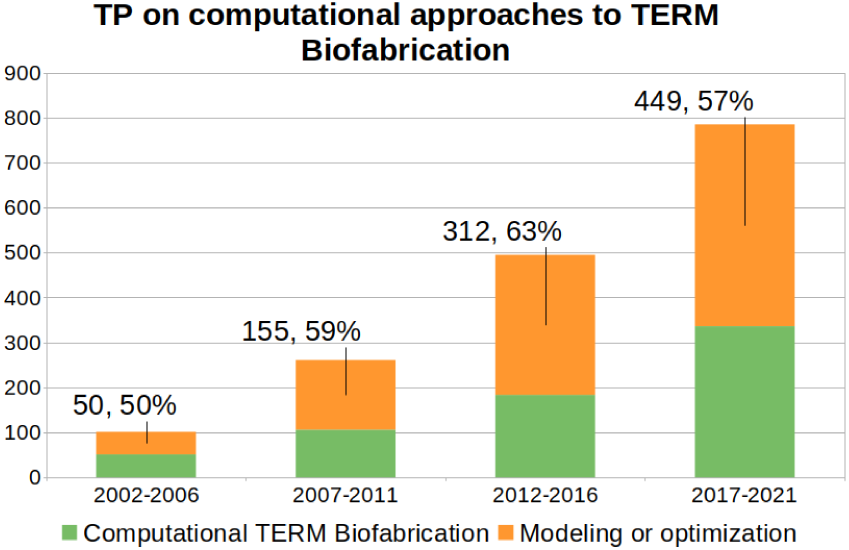
Total Publications on *Computational approaches to TERM Biofabrication* (*Query 2.1*) aggregated over five-years periods, with a highlight on the contributions on *Computational approaches to TERM Biofabrication based on either modeling or optimization* (*Query 2.2*), that constitute 50% of TP between 2002 and 2007 (50 publications), grow to 63% for the period between 2012 and 2016 (312 publications), and decrease to 57% for the period between 2017 and 2021 (449 publications).

**Figure 6:**
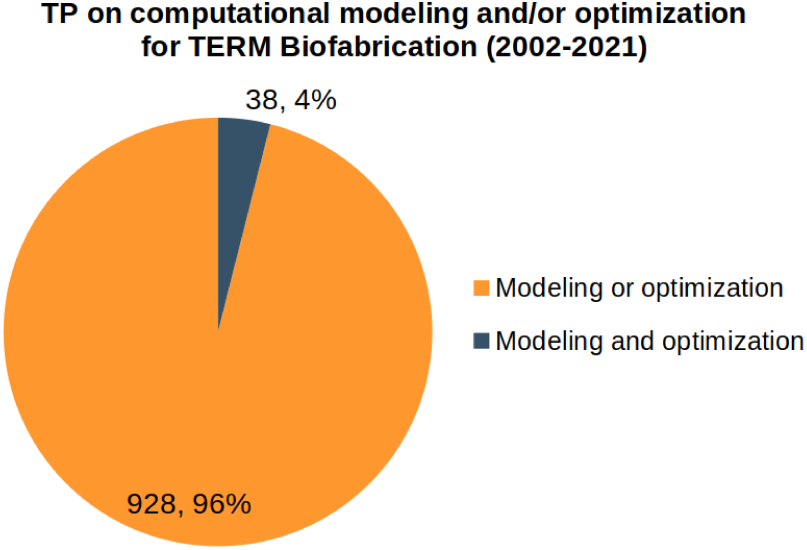
Total Publications between 2002 and 2021 on *Computational approaches to TERM Biofabrication based on either modeling or optimization* (*Query 2.2*, 966 TP), highlighting the 4% (38 TP) on *Computational approaches to TERM Biofabrication based on combined modeling and optimization* (*Query 2.3*).

### 2.2. Science Mapping

After quantitatively analyzing publications in the reference scientific domains, this section aims to analyze and visualize the structure of these domains by performing science mapping [13]. The analysis leveraged the VosViewer tool [14, 15] to search publications on the PubMed platform [16]. The analysis focused on results of Queries 2.2-3 summarized in Figure 5 and Figure 6.

The first step of the analysis was constructing a cooccurrence network from the PubMed file generated with Query 2.2. The co-occurrence network emerged from text analysis of the *Title* and *Abstract* fields of each publication, then performing clustering analysis over the obtained network. Due to the high heterogeneity of terms in text data, this step employed a thesaurus file. Different versions of the same terms or concepts (plurals or singulars, synonyms, very close terms) were considered under the same umbrella term to facilitate network homogeneity and visualization.

Figure 7 shows the text-based co-occurrence map based on Query 2.2. Different colors indicate separate clusters within the network. While the green cluster at the top includes clinical and biological concepts, the blue cluster at the bottom centers on technological approaches and research organization, and the smaller third cluster in orange contains terms related to biomaterial properties. Interestingly, terms about computational approaches exist in both the blue and orange clusters, with a predominance in the larger blue cluster. Figure 8 provides an enlarged view over the portion of the map where terms referring to computational approaches emerge.

**Figure 7:**
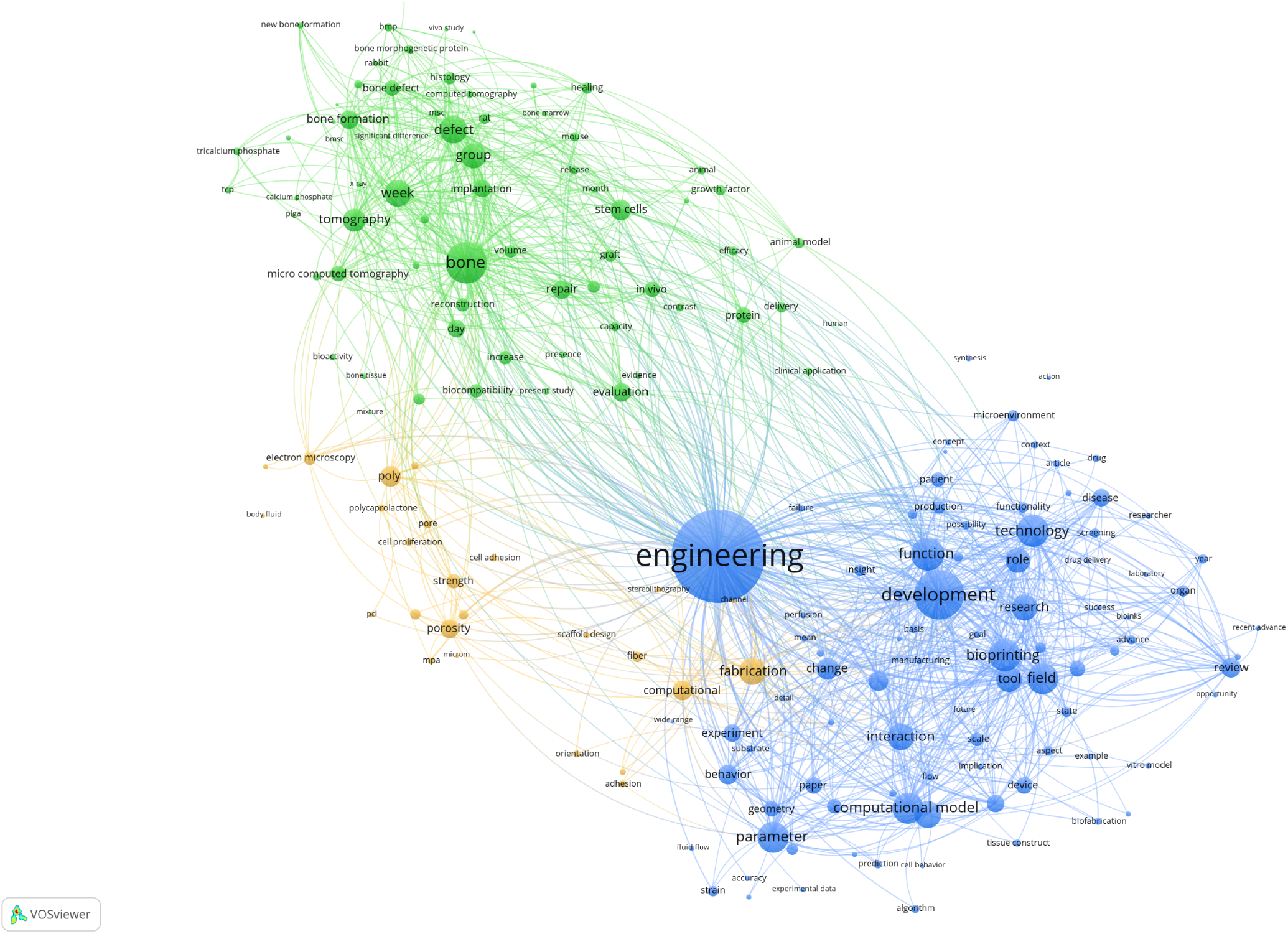
Text-based terms co-occurrence map of *Computational approaches to TERM Biofabrication based on either modeling or optimization* (*Query 2.2*). The map is based on text analysis over PubMed files and extracts co-occurrence information from the bibliographic database file obtained from the PubMed platform with *Query 2.2* performing binary counting of terms occurrences over the *Title* and *Abstract* fields, and considering the 60% most relevant terms among those occurring in at least ten documents (290 over 483). The relative size of term nodes indicates total occurrences and link strength indicates co-occurrences. Colors indicate the three different clusters in the network. Clustering relied on Association Strength as a normalization method and a Resolution parameter value of 1.00.

**Figure 8:**
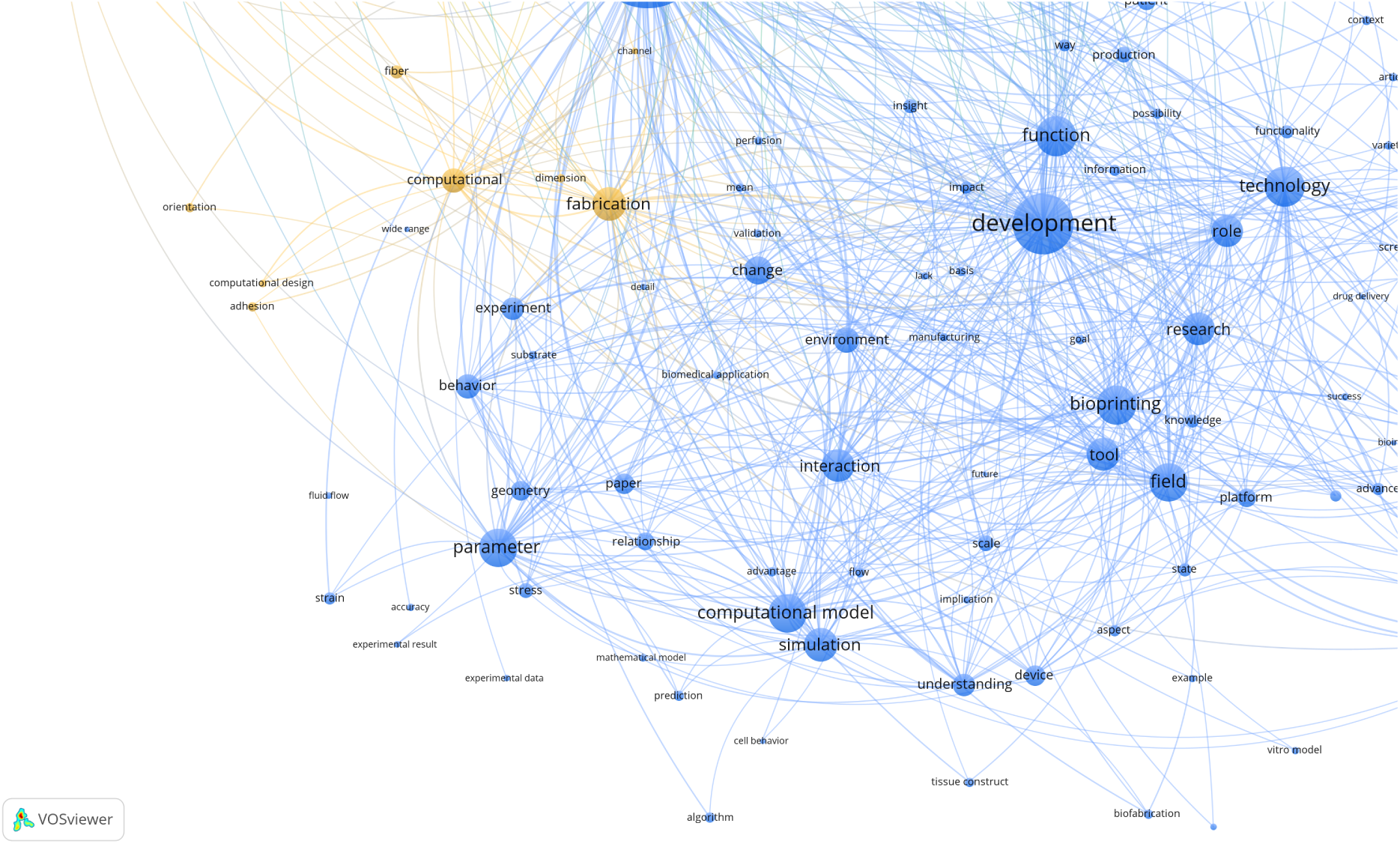
Enlarged portion of the text-based terms co-occurrence map of Total Publications between 2002 and 2021 on *Computational approaches to TERM Biofabrication based on either modeling or optimization* (*Query 2.2*). The map is based on text analysis over PubMed files. It extracts co-occurrence information from the bibliographic database file obtained from the PubMed platform with *Query 2.2* performing binary counting of terms occurrences over the Title and Abstract fields and considering the 60% most relevant terms among those occurring in at least ten documents (290 over 483). The relative size of nodes indicates term occurrences, and link strength indicates co-occurrences. Colors indicate the three different clusters in the network. Clustering relied on Association Strength as a normalization method and a Resolution parameter value of 1.00.

A simplified text-based co-occurrence map on Query 2.2 devising a different thesaurus file was also created to simplify situations where different concepts of the same domain group under the same umbrella terms. In particular, all terms related to modeling, simulation, and computational aids combine under the term “computational approaches." “Regenerative medicine," “scaffolds," and “bioprinting" similarly grouped several synonyms and adjacent terms. This approach created nodes with higher total occurrences and aggregated co-occurrence links, creating a simpler network that was easier to visualize.

Figure 9 shows the resulting simplified co-occurrence map, which has two clusters. The green cluster on the bottom left centers over biological and clinical aspects. In contrast, the blue cluster on the top right centers on technology. Computational approaches belong to this cluster.

**Figure 9:**
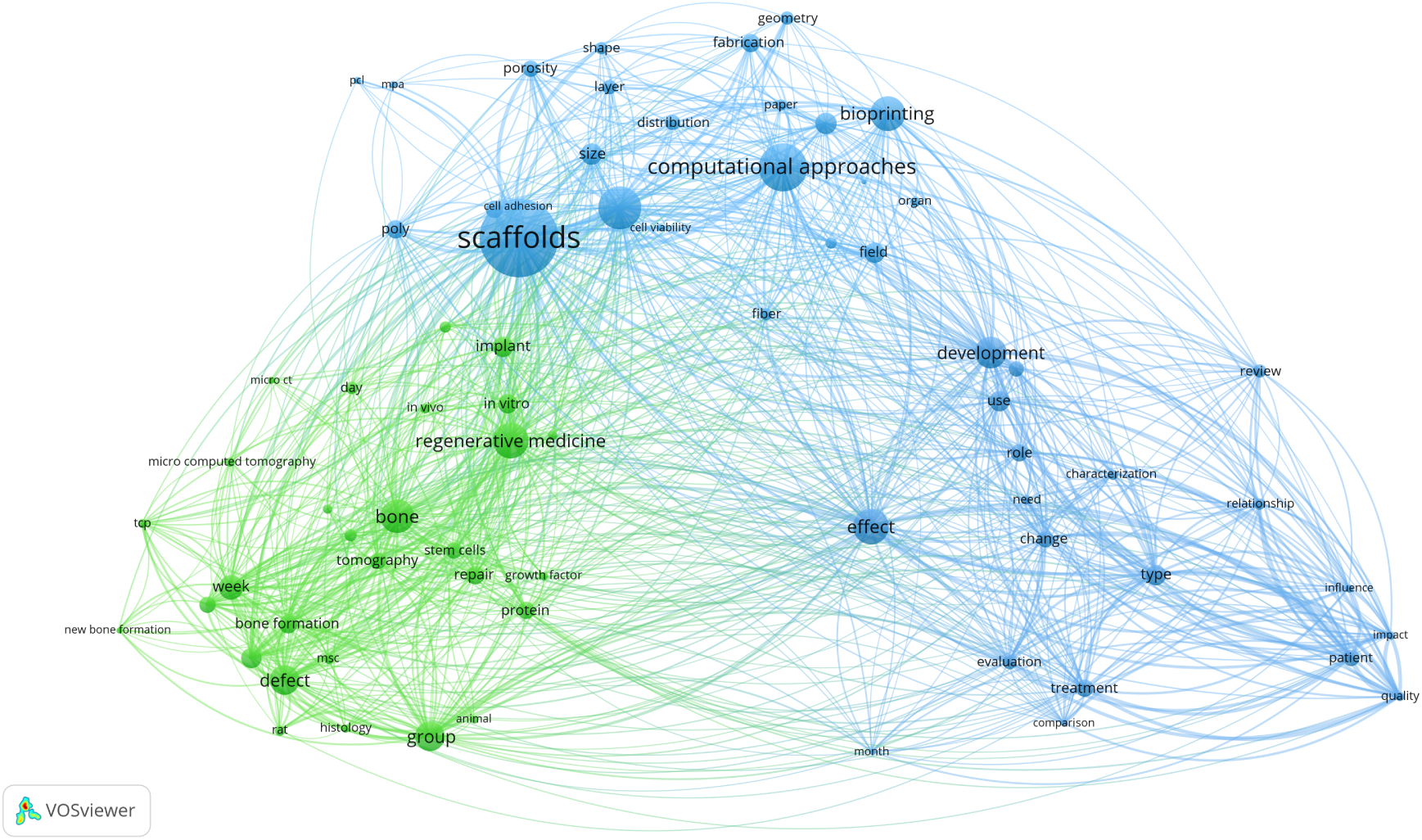
Simplified text-based terms co-occurrence map of *Computational approaches to TERM Biofabrication based on either modeling or optimization* (*Query 2.2*). The map is based on text analysis over PubMed files and extracts co-occurrence information from the bibliographic database file obtained from the PubMed platform with *Query 2.2* performing binary counting of terms occurrences over the *Title* and *Abstract* fields, and considering the 60% most relevant terms among those occurring in at least 30 documents (151 over 24820). The relative size of term nodes indicates total occurrences and link strength indicates co-occurrences. Colors indicate the three different clusters in the network. Clustering relied on Association Strength as a normalization method and a Resolution parameter value of 1.00.

The second step of this analysis devised a second map from bibliographic data summarizing results of Query 2.2, in particular creating a co-occurrence network of MeSH keywords, employing the Fractional counting method, and considering only the 38 keywords occurring at least three times. Due to the high homogeneity of MeSH keywords in bibliometric data, this step did not employ a thesaurus file. Figure 10 illustrates the co-occurrence network of keywords obtained and the two clusters identified. The top blue cluster includes more details about the biological aspects of tissue engineering. The green cluster on the bottom contains more terms about computational modeling and analysis approaches.

**Figure 10:**
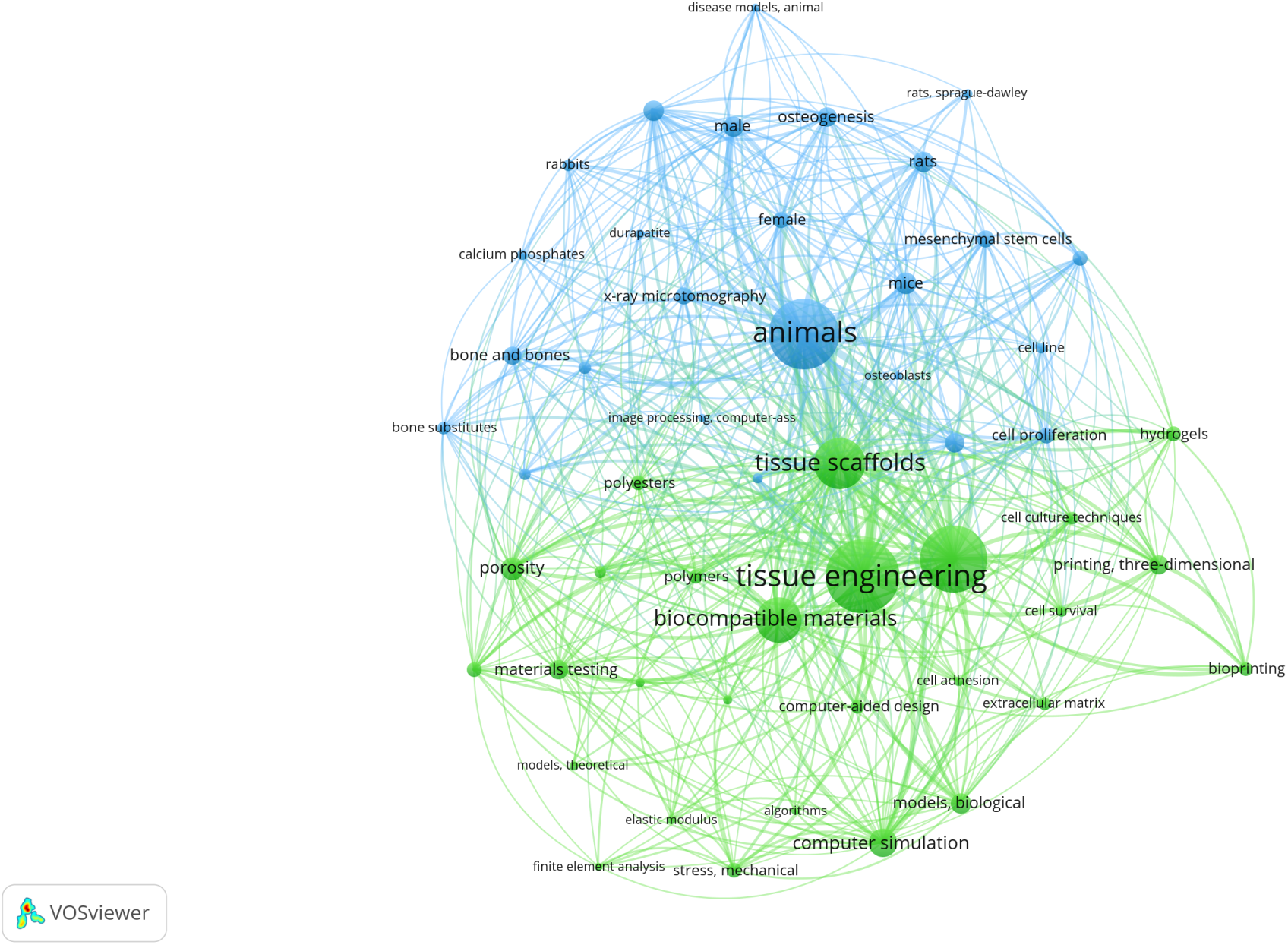
Bibliographic data-based co-occurrence map of *Computational approaches to TERM Biofabrication based on either modeling or optimization* (*Query 2.2*). The map is based on a bibliographic analysis of PubMed files. It extracts MeSH keywords co-occurrence information from the bibliographic database file obtained from the PubMed platform with *Query 2.2*, performing Fractional counting of keyword occurrences and considering the ones occurring in at least 30 documents (52 over 1348). The relative size of nodes indicates keyword occurrences and link strength indicates the number of co-occurrences. Colors indicate the two different clusters in the network. Clustering relied on Association Strength as a normalization method and a Resolution parameter value of 0.8.

To conclude, the analysis targeted the subset of 38 TP obtained with Query 2.3, and Figure 11 shows the text-based terms co-occurrence network with the related clusters.

**Figure 11:**
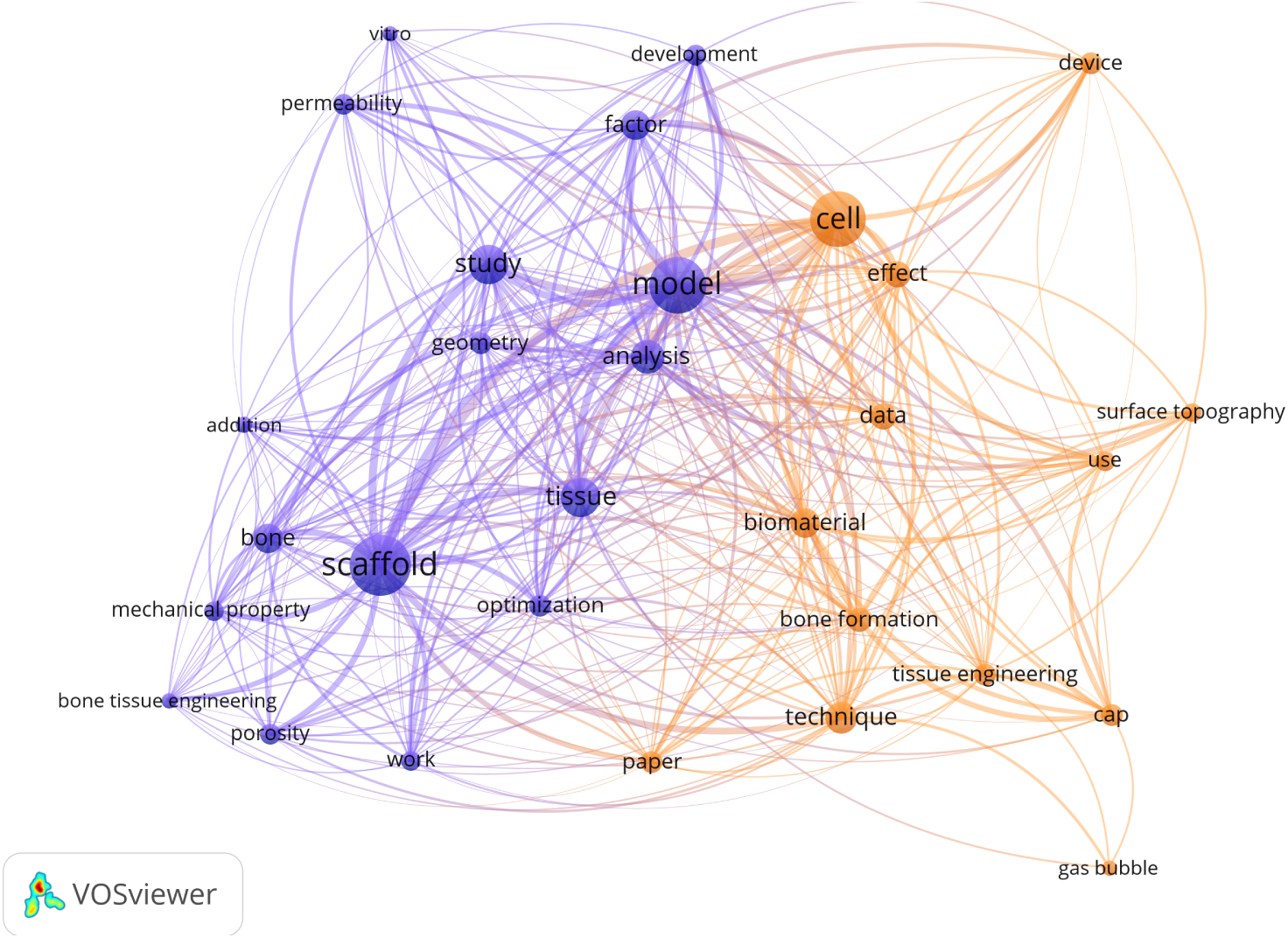
Text-based terms co-occurrence map of *Computational approaches to TERM Biofabrication based on modeling and optimization combined* (*Query 2.3*). The map is based on text analysis over PubMed files. It extracts co-occurrence information from the bibliographic database file obtained from the PubMed platform with Query 2.3 performing full counting of terms occurrences over the *Title* and *Abstract* fields and considering the 30 terms occurring in at least six documents. The relative size of term nodes indicates total occurrences and link strength indicates co-occurrences. Colors indicate the three different clusters in the network. Clustering relied on Association Strength as a normalization method and a Resolution parameter value 0.70.

### 2.3. The untapped potential of computational methods for TERM biofabrication

Together, the performance analysis and science mapping results in this section suggest that the scientific interest in TERM biofabrication is rapidly growing. Publications on computational approaches to TERM biofabrication constitute a small but constant portion of scientific production, and about half of them consistently center on modeling or optimizing biofabrication processes. Approaches that combine simulation and optimization are a small subset of the total publications on either simulation or optimization: current approaches tackle specific computational problems separately, in addition to separate biofabrication process stages (see Section 4). Science mapping shows that computational approaches link several clinical and biological aspects and biofabrication technologies. These findings highlight the applicability of computational approaches to many aspects of biofabrication and the lack of comprehensive approaches targeting multiple of its challenges together, describing their untapped potential to tackle different stages and issues in designing a biofabrication process.

## 3. Computational methods for intelligent biofabrication: the technological context

In order to provide a context to this review, the following sections analyze the significant innovative trends in TERM biofabrication that are paving the way for extensive usage of computational methods in this field. In particular, they highlight the growing tendency of combining multiple technologies in biofabrication (subsection 3.1), the increasing diffusion of automation and digitalization to optimize bio-fabrication processes (subsection 3.2), the employment of research design techniques (subsection 3.3) and finally the importance of comprising process complexity into computational approaches to biofabrication (subsection 3.4).

### 3.1. Multi-technology biofabrication harmonizes combined capabilities

Biomimetic TERM products must replicate the complex and hierarchical structure of their *in vivo* counterparts. However, individual biofabrication technologies aim to control particular aspects of the final product. Such limitation appears even when considering a single class of technologies. Biomimetic scaffolds, for instance, must have a hierarchical architecture, including application-specific surface properties. However, each fabrication approach usually targets a specific resolution range, which limits fabricating hierarchical structures [2]. In order to overcome this limitation, hybrid biofabrication process schemes combine different fabrication approaches to expand the dimensional range covered and thus achieve biomimetic architectures [17].

Extending this reasoning to different classes of technologies, one can say that biofabrication approaches are today designed to control partial aspects of the final product. For example, bioprinting approaches control spatial organization. Automated culture systems control functional maturation. Directed differentiation of cells governs the functional specialization of the biological building blocks for biofabrication.

Therefore, to control product quality, biofabrication processes can combine different technologies. Multi-technology biofabrication defines the combination of heterogeneous technological approaches, ranging from additive manufacturing to automation and computational process design, working together towards fully biomimetic TERM products [6]. Multi-technology biofabrication goes beyond combining different techniques, focusing more on their synergistic integration. Computational methods are crucial in enabling integration and rational harmonization in multi-technology procedures.

### 3.2. Automation and digitalization optimize processes

Biomanufacturing is moving towards fully integrating the Industry 4.0 principles in designing, executing, and optimizing manufacturing processes. Indeed, research automation and digitalization are spreading in the life sciences domain. For instance, fast biofoundries in synthetic biology are *“specialized laboratories that combine software-based design and automated or semi-automated pipelines to build and test genetic devices"* [18].

Several biomanufacturing applications rely on Robotic Process Automation (RPA), i.e., the use of a *“pre-configured software instance that uses business rules and predefined activity choreography to complete the autonomous execution of a combination of processes, activities, transactions, and tasks […]"* [19, 20]. Digitalization is a fundamental part of process automation, and it includes two main aspects: the *digitization* of existing operations and the creation of *digital twins* of existing processes. The *digitization* of existing laboratory processes is the transformation of analogical and manual operations into digital and semi-automated processes, i.e., the compilation of a laboratory diary or the acquisition and storage of newly collected data. Digitization and automation can work together to increase biofabrication process traceability, control, and quality [21].

*Digital Twins (DTs)* [22, 23], are *“complete virtual descriptions"* of a physical process that are *“accurate to both micro and macro-level"* (adapted from [24]). They act as a digital counterpart of a physical process, dynamically modeling and analyzing it to actuate and modify the system in a risk-free environment, supporting new understanding and rational organizing of research activities. These technologies can serve different scopes in biomanufacturing, from version control systems for synthetic biology [25] to bioprocess modeling [26] and optimization [27]. DTs and computational approaches have a growing role in developing biomimetic TERM biofabrication products by supporting process execution optimization and intelligent process design [28, 29].

### 3.3. Research design goes beyond empirical trials

Empirical methods face inherent limitations in exploring extensive and complex process design spaces due to the high resource costs and time demands associated with *in vitro* experiments. Automation and digitalization can enhance *in vitro* experimental campaigns by enabling a more significant number of experiments, thus broadening the exploration of the design space. However, adhering strictly to a trial-anderror experimental paradigm, even with these advancements, often leads to only incremental innovations, improving existing process designs based on literature that might be incomplete, unreliable, and non-reproducible [30], and often chosen for popularity rather than performance [31].

Traditional trial-and-error methods, such as One Factor at A Time (OFAT) campaigns, that explore ranges of relevant system parameters one at a time [32], are costly in terms of time and resources. They also fail to recognize interdependencies among system variables, which hampers the ability to connect experimental results to realistic process designs. Actual designs control multiple variables simultaneously. These limitations result in OFAT yielding suboptimal processes and products [23].

In contrast, factorial experimental design, which supports testing under multiple conditions, has proven effective in investigating optimal conditions for stem cell differentiation [33]. Design-of-Experiments (DoE) computational techniques [34, 23] enable a more efficient, systematic exploration and exploitation of complex design spaces [32, 8, 35], showing adequacy in tackling multi-factorial problems in the optimization of directed cell differentiation [36, 37] and in the development of tissue engineering scaffolds [38].

### 3.4. White-box models to optimize process design

Artificial Intelligence (AI) technologies offer the potential to automatically adjust experimental strategies as new data is generated, thereby maximizing information extraction and enhancing process improvement efficiency [7, 39].

Computational tools and AI are instrumental in moving beyond traditional experimental trial-and-error methods [40, 41], paving the way for the adoption of intelligent automation, which utilizes AI in automated and digitized processes to aid research design and result analysis [40, 41].

Computational Design Space Exploration (DSE) benefits from the integration of computational biofabrication models with optimization strategies, enhancing both research and process design by increasing the accuracy of bioprocess representation [34] and applying it for process optimization [7, 39]. The design space of a biomanufacturing process is a multidimensional space defined by input variables and process parameters that influence product quality [5]. However, modeling the intricate biological complexity inherent in biofabrication processes presents substantial challenges to model-based DSE approaches.

In this direction, several computational approaches sustain biofabrication in general [29, 42], and its process modeling in particular. While Machine Learning (ML) and Artificial Neural Networkss (ANNs) offer black-box models of the system, computer simulations provide white-box models that capture mechanistic relationships and analyze complex dynamics under various conditions [43]. To support model-based DSE, white-box models of biofabrication must be accurate and predictive, as well as strike a balance between the holistic and hypothesis-driven modeling approaches.

Holistic modeling takes a comprehensive approach, encompassing numerous components and interactions, akin to viewing the entire system through a wide-angle lens. These models need to consider multi-level systems, from individual bioprocesses to multicellular aggregates [44], and are expected to support dynamic simulations of complex biological processes [45, 46]. Although excellent for identifying emergent properties, the complexity of such models can be computationally intensive and come with many uncertain parameters [47]. High model complexity creates a vast and probabilistic design space, challenging both *in vitro* experimentation [48] and computational DSE.

Conversely, hypothesis-driven modeling focuses on testing specific aspects of the system [49], offering more targeted and manageable studies but potentially overlooking broader system interactions. This approach yields simpler models with lower computational demands, facilitating model-based computational DSE.

Ultimately, holistic and hypothesis-driven models respond to the challenge of incorporating biological complexity into biofabrication models. Both approaches can include the biological aspects of biofabrication processes to support their computational designs. Thus, both approaches or a combination thereof in hybrid models [50] provide valuable white-box modeling tools [49], with a positive impact on TERM biofabrication.

## 4. Computational methods for intelligent biofabrication: a review of state-of-the-art solutions

As thoroughly reviewed in [51], process design, modeling, and optimization find many applications to the different stages of TERM biofabrication. This section provides a review of the state-of-the-art computational methods applied to TERM biofabrication, including and not limited to the publications from Query 2.3 (see Section 2), focusing on techniques combining modeling and optimization methods. The review organizes around selected critical stages for TERM biofabrication [52], in particular: *Product modeling and quality control* (Figure 12.A-E, subsection 4.1), *Biomaterials qualification* (Figure 12.B, subsection 4.2), *Fabrication* (Figure 12.C, subsection 4.3) and *Maturation* (Figure 12.D, subsection 4.4).

**Figure 12:**
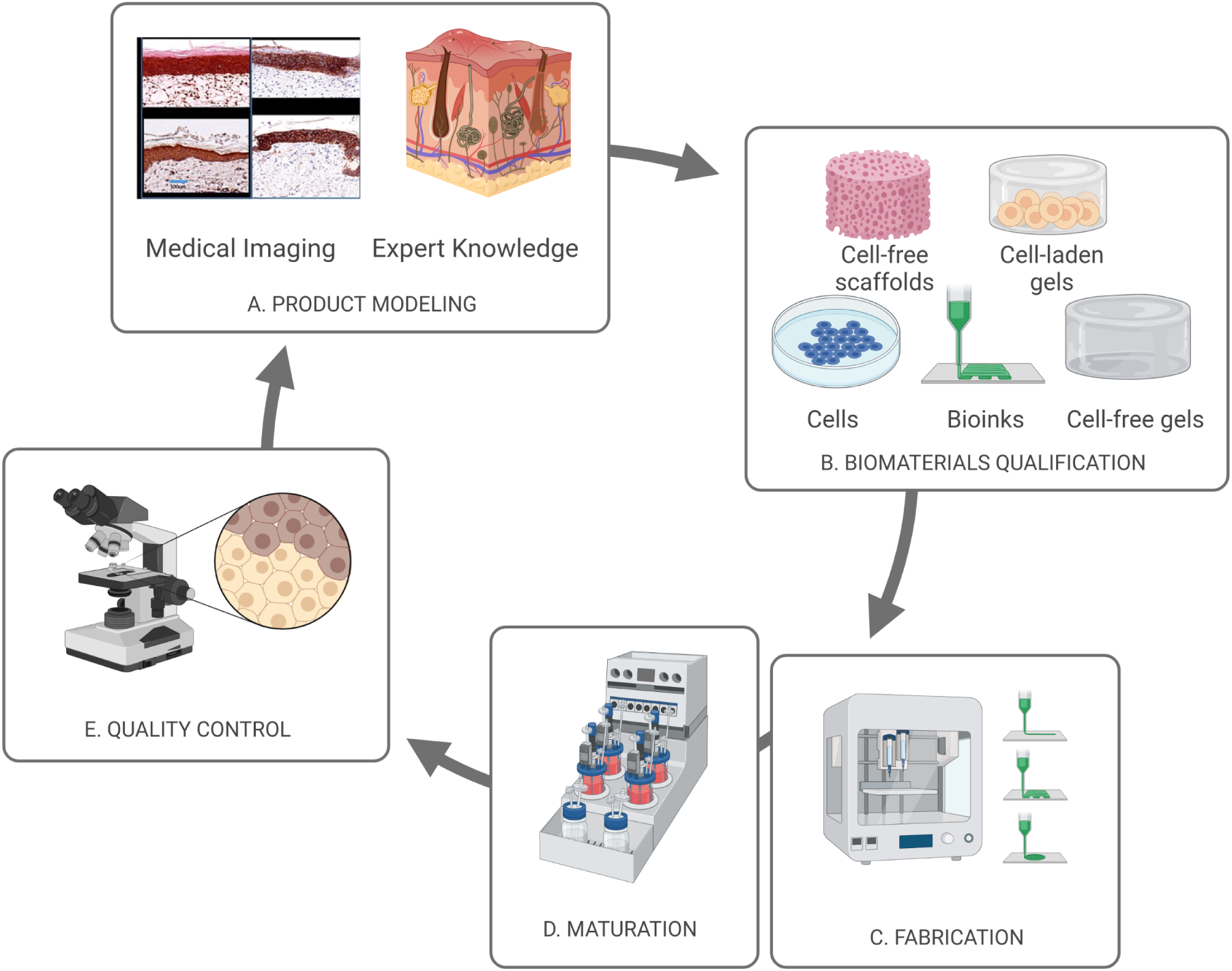
Overview of a biofabrication process: (A) medical images and expert knowledge define a model for the target product, which guides the (B) choice and combination of biomaterials and cells used for (C) fabrication via bioprinting and bioassembly, followed by (C) construct maturation; (D) evaluation of the quality of the mature product relies on measuring its closeness to the structural and functional features of the defined target.

### 4.1. Product modeling and quality control

Product modeling (Figure 12.A) aims at defining the target product from expert knowledge or following a datadriven process using medical imaging to determine the desired features in the product. Quality control (Figure 12.E) assesses the presence of the desired functional and structural features defined in the product model, especially if biofabrication has a biomimetic purpose (Figure 12.A). Product modeling includes retrieving and using information on the product, its design, and the definition of the desired features. The ultimate goal of TERM biofabrication is fulfilling a patient’s clinical need for regeneration. This goal implies that a biofabricated product must be biomimetic; that is, it must faithfully recapitulate the biological structures to regenerate. This principle guides the product design and modeling and defines quality metrics for the biofabrication products obtained. In particular, for TERM applications, the quality of the product is often reflected in biomimetic fidelity to its *in vivo* counterpart since the ultimate goal is regeneration.

#### 4.1.1. Data-driven reconstructions

Computational tools support automated data-driven modeling of the functional and structural features to replicate the natural system, increasing biomimetic fidelity in a TERM biofabrication product. Image-based Computer-Aided Design (CAD) models can build on processed images acquired directly from patients, including Magnetic Resonance Imaging (MRI) or Computed Tomography (CT) scans that are later converted to Digital Imaging and Communications in Medicine (DICOM) format [53] and then to Standard Tassellation Language (STL) files for printing [54]. This process finds application in orthodontic regeneration [55] and soft-tissue engineering [56] as well as in the generation of three-dimensional (3D) heart valves [57].

However, tissue images *per se* fail to provide information at the single-cell level. Thus, reconstructed 3D tissue models usually lack multi-scale cellular resolution and tissue-or organ-specific properties.

Computational methods support the development of TERM product models beyond image processing, by allowing the discovery of tissue architectures from imaging and spectral data, bridging the information from the *in vivo* tissue microenvironment to the construct properties at the scale that the bioprinting technology chosen can fabricate [58]. To this aim, Deep learning (DL) super-resolution unsupervised approaches [59] can transform low-resolution images into higher-resolution versions, narrowing the gap between the average resolution of medical images on the millimeter scale and the resolution of bioprinting technologies, which lies on the micrometer scale [60]. This transformation allows extracting biological features at the cellular level in an unbiased and data-driven manner. For example, authors in [61] leveraged a Convolutional Neural Network (CNN) to classify cells according to their cell cycle phase. These initiatives support the data-driven construction and enrichment of Intelligent Digital Twins (IDTs) comprising a multi-scale model of the target products’ architectural and functional features, for which is crucial to build and curate supporting databases [62].

#### 4.1.2. Computer-assisted design

CAD is a technology used for creating precise two-dimensional (2D) and 3D models of physical components, widely used in industries such as engineering, architecture, and manufacturing. It improves design accuracy and efficiency and facilitates easy modifications, often integrating with other technologies like Computer-Aided Manufacturing (CAM) and 3D printing for simplified product development. Bio-CAD, bio-CAM, and bio-CAE apply CAD, CAM, and Computer-Aided Engineering (CAE) to biological processes and subprocesses. The bio-CAD process supports the design of tissue and organ blueprints, the bio-CAM method the manufacturing biological products, and the bio-CAE the creation of complex architectures, and the validation and optimization of biomanufacturing tools and bioproducts [63, 64]. CAD is the most widely employed method for designing scaffolds from scratch or using 3D scans of the target biofabrication product [65], and supports the design of scaffolds providing the localized control of biomolecules distribution for tissue engineering and drug release [66].

Implementing bio-CAD models also relies on computational methods. Several approaches exist to translate a model into a physiologically relevant product through computer-controlled Rapid Prototyping (RP) [67, 68, 69]. Computational modeling of tissue- and organ-level biological complexity combine with bio-CAD to embed complex functionality into TERM products, optimizing them and increasing their quality. For instance, ML techniques predict material properties related to various mixture compositions of the bio-inks and support scaffold designs by learning from an extensive database of materials and designs [70, 71]. ML also supports multi-objective optimization of material process variables [72].

Biofabrication products are poised for subsequent maturation, adapting to the stimuli received during maturation. Thus, product design must consider the cellular component’s role in the product’s evolution to its final form, requiring consideration of the product’s evolution after the initial construction and under different environments. Computational approaches support this by guiding product design considering potential functional responses to different stimuli [73], following the four-dimensional (4D) printing paradigm [74], producing dynamic structures whose functionality, shapes, and properties change based on the environmental inputs. Since cells are the main actors of transformation and response to the environment, product design must consider their behavior in predicting the final product form. Computational tools help in this by modeling intracellular, cellcell, and cell-biomaterials interactions during fabrication [62] and subsequent maturation [75, 76, 77].

This concept finds unique declination according to the specific biofabrication product and regenerative application. For example, integrating computational modeling in heart valve design could predict long-term in vivo performance, remodeling, and failure under non-physiological pressure loading in a translational sheep model [78]. Bioprosthetic heart valve (BHV) presents unique challenges for computational design: standard Finite Elements Methods (FEM) can simulate the effect of hydrostatic forcing on a closed BHV, but this approach fails to account for transient responses during valve opening and closing. Nonetheless, the latter effects are not negligible since they contribute to long-term structural fatigue and thus to the risk of BHV transplant failure. Thus, BHV computational design must rely on models of the surrounding hemodynamics that support the accurate simulation of these effects, as provided in [79], where authors propose a complete mechanical model of a BHV based on the immersogeometric Fluid-Structure Interaction (FSI) methodology.

In conclusion, computational methods support the product design and quality in TERM biofabrication by automated data-driven extraction of target product features and by handling the complex computer-assisted design of dynamic products that evolve independently and after complex responses to environmental stimuli.

#### 4.1.3. Scaffolds

Porous scaffolds for TERM products must have certain biological, biomechanical, and biomaterial-related functionalities, including suitable pores size and geometry, allowing nutrients to diffuse uniformly and cells of the engineered tissue to invade the scaffold. Secondly, scaffold surfaces must have physical-chemical properties, ensuring biocompatibility; that is, they must allow cells to adhere and develop the expected phenotypes. During fabrication, scaffold material must exhibit mechanical properties supporting the bearing of required loads during regeneration. Finally, scaffolds must be biodegradable or bioabsorbable once sufficient tissue has formed to guarantee mechanical support to the TERM product.

The role of computational methods in scaffold fabrication is to identify optimal trade-offs considering this set of complex requirements [80, 38]. Moreover, the comprehensive analytical and numerical modeling of the 3D and 4D additive manufacturing benefits biofabrication with understanding, predictive modeling, and optimization of the biofabrication process at different scales [81].

Most computational approaches in the literature target optimizing the mechanical and chemical properties of bone graft scaffolds [82], including porosity, micro-architecture, Young’s modulus, and dissolution rate. In [83], the authors propose a general design optimization scheme for 3D internal scaffold architecture to find a good trade-off between elastic properties and porosity. They introduce the homogenization-based topology optimization algorithm, demonstrating that the method can produce biomimetic structures mimicking anisotropic bone stiffness obtained with scaffolds of widely different porosity. In [84], authors propose a 3D computational model based on Sussman–Bathe hyperelastic material behavior to study interpenetrated polymer networksfor cartilage repairing, quantitatively simulating the distribution of fluid fluxes and nutrient supply within the different regions of a construct.

While these properties are pivotal for scaffold quality, considering cell-material interactions is crucial to the modeling and designing of TERM bone grafts. Computational design must account for the influence of growth factors and other biochemical signals and the accurate simulation of dissolution processes, including the impact of degradation products such as calcium ions and inorganic phosphate on bone formation biological processes. In bone tissue engineering, the passage from scaffold design to manufacturing requires compliance with the design control requirement that must undergo verification in a finished device. Authors in [85] propose a computational approach to investigate the accuracy of the final properties of polycaprolactone scaffolds fabricated by selective laser sintering [86].

On the structural optimization front, Topology Optimization (TO) aims to optimize scaffold printing by organizing material placement in space. For instance, different TO approaches such as Solid Isotropic Material with Penalization (SIMP) [87], Bidirectional Evolutionary Structural Optimization (BESO) [88], and level-set [89] support the design of differentially dense and stiff structures. In [90], authors leverage TO on the performance of a chitosan hydrogel bioprinted soft actuator while maintaining the material’s volume fraction. Triply periodic minimal surfaces (TPMS) recently inspired the fabrication of biomimetic porous scaffolds since they support cell adhesion, migration, and proliferation. A versatile design method for TPMS sheet scaffolds, which can satisfy multiple requirements simultaneously, was proposed in [91]. In these applications, TO does not consider cell-related aspects: optimized parameters only involve a single scale and cell-free biomaterials.

However, the geometric properties of scaffolds for tissue engineering directly affect cellular deposition. Computational tools sustain the design of scaffolds with varying controlled geometrical and functional properties throughout their structure [92, 93], such as varying porosity [94]. Models of cellular behavior can help choose geometrical properties ranges to use for scaffolds. For example, computational models of osteoblasts support the prediction of the growth of bone matrix tissue in scaffolds with different pores geometry [95].

Nevertheless, TO approaches easily extend to cell-laden and multi-scale biomaterials. For instance, in [96], authors present a level-set-based topology optimization algorithm and a time-dependent shape derivative to optimize scaffold architecture for femur models, showing significant advantages in continuing bone growth compared to stiffness-based topology optimization, time-independent design, and typical scaffold constructs. The common usage of topology optimization and models of biological responses to different scaffold properties prove effective in supporting scaffold design [97]. In [98], authors leverage Three Dimensional Convolutional Neural Network (3D-CNN) surrogate models to optimize multi-scale topology on non-parametric microscale structures. Authors in [99] present different computational approaches, including surface hit detection and ordering and prediction of relevant surface properties and cell responses, aiming at optimizing surfaces to influence cell behavior.

In this direction, FEM is a powerful modeling and optimization tool for biomechanical research [100, 101, 102, 103, 104], which supports the modeling of not only biomechanical and geometrical aspects of the cell-free scaffold, but also of dynamical features and the responses of specific cell types during later construct maturation [105]. Authors in [106] optimize geometrical features of scaffolds, maximizing bone growth rate with an algorithm relying on parametric Finite Elements (FE) models of scaffolds, computational models of cells, and optimization methods.

Similarly, reaction-diffusion models [107] support the evaluation of geometric and mechanical properties of scaffolds [108], as well as the rational design of waveguides for cells within biomaterials [109]. The computational simulation of 3D bone tissue regeneration based on the voxel FE method, including new bone formation and scaffold degradation, is at the core a model-driven computational framework for designing and optimizing a porous scaffold microstructure in [110].

Computational modeling of the bone formation process helps design optimal combinations of calcium-based biomaterials and cell culture conditions to maximize the quantity of formed bone in [111]. FE analysis, Computational Fluid Dynamics (CFD) and mathematical modeling support the choice of optimal mechanical properties of a scaffold for cartilage regeneration [112, 113]. The simulation of matrix degradation processes informs the fabrication of a cartilage-like construct to study cartilage degradation in osteoarthritis in [114].

Considering cardiovascular applications, a notable example is scaffolds for *in situ* cardiovascular tissue engineering. This approach uses a synthetic, biodegradable scaffold as a temporary framework for incoming cells to develop their own Extra-Cellular Matrix (ECM), which gradually breaks down, allowing tissue regeneration. Computational models are pivotal in factoring in the role of incoming cells in product quality, directly impacting the clinical outcome [115].

Vascularization of constructs is still a major limiting factor in the size and quality of TERM products. Computational design must comprise the multiple interdependent regulatory mechanisms in the cross-talk between endothelial cells and native tissue cells to recapitulate the formation of new functional blood vessels. A computational model of vascular adaptation and a formal optimization method support tissue-engineered vascular graft design in [116]. Computational simulations and microvascular network analysis supported optimization of the geometry and oxygen distribution within hydrogel constructs in [117].

Scaffolds are crucial in fabricating biomimetic liver models [118, 119]. Computational modeling finds application in the characterization of vascular load in scaffolds for liver engineering for their rational design [120, 121].

For wound healing applications, computational tools support the image-based automatic structural design of scaffolds and localized control of biomolecule distribution to sustain healing over time [66, 122].

Another example of the unique challenges posed to scaffold design by different TERM applications is nerve regeneration. The computational design allowed exploring the impact of various structural features on their properties to increase biomimicry and optimize porosity and permeability for the design of nerve guidance conduits, tubular tissue engineering scaffolds used for nerve regeneration [123].

In conclusion, computational methods support scaffold design and optimization in TERM biofabrication by automatically finding optimal trade-offs between application-specific requirements, which also extends to the requirements posed by the complex cellular processes they host.

### 4.2. Biomaterials qualification

The definition of the product model guides the selection of biomaterials and cells to employ (Figure 12.B). TERM biofabrication relies on either the bioassembly of building blocks based on living cells, bioprinting of biomaterials, or a combination of the two [124] (see subsection 4.3). Therefore, the qualification of inorganic and organic biomaterials used for these two processes dramatically affects the quality of the final product. The specific combinations of biomaterials must carefully adjust to the scopes and machinery available for biofabrication. For example, a bioink must be biocompatible with the cell type to be printed and printable using bioprinting technology. Cells for preparing cell-laden materials or seeded in cell-free scaffolds undergo selection, culture, and differentiation before fabrication. Computational tools sustain such biomaterials’ rational design and development in several ways, combining the non-trivial composition of multiple mechanical and biological properties, which often conflict.

#### 4.2.1. Inorganic biomaterials

Computational approaches support biomaterials characterization and explore optimal trade-offs between different desired features of biomaterials. A notable example in bioprinting is the trade-off between the degree of biocompatibility for the cells employed and the printability, considering the bioprinting technology of choice. Failure Modes and Effects Analysis (FMEA) and Quality Function Deployment (QFD) support the identification of functional requirements for a specific application in the design of a new biomaterial. In contrast, Multi-attribute Decision Making (MADM) approaches support selecting the best constituent materials to employ. The combination with DoE allows searching the biomaterial design space efficiently, saving on experimental resources [125, 126].

Extending these approaches, Multi-objective Decision Making (MODM) tackles biomaterial design problems that seek to satisfy multiple goals concurrently, e.g., adequate stiffness and degradation properties in bone scaffolds.

Computational multi-objective optimization provides multiple optimal solutions. Computational models of biomaterials support DSE by testing the optimal solutions *in silico*. For example, a model of viscoelastic materials mimicking soft biological tissues demonstrates its capability to capture complex viscoelastic behavior observed experimentally [127]. Such accurate, validated models of physical and chemical biomaterial behaviors support the computational testing of the generated optimal solutions to prioritize subsequent experimental tests further, saving time and resources.

This reasoning extends to cell-laden biomaterials. Embedding computational models of biological components into biomaterial design and optimization for the design of biomaterials, providing more accurate and comprehensive product quality prediction, further minimizes the experimental costs of iterative material synthesis and testing [38]. In this direction, modeling of organic and inorganic components, combined with optimization solutions, supports the characterization and the choice of optimal biomaterials composition given a specific TERM application. For example, FE modeling and optimization support the characterization of polyvinyl alcohol as a potential biomaterial for cartilage tissue engineering scaffolds in [128]. The joint consideration of both inorganic substrates and cellular components allowed the creation of an *in silico* design library based on numerical modeling to predict composite biomaterials performance and support their design, applying it to articular cartilage engineering for patients in [129].

#### 4.2.2. Organic biomaterials

While several computational approaches target the design and optimization of inorganic biomaterials, factoring in their interactions with cells, other approaches focus on the cellular component only, in particular, to guide the selection, design, and culture of cells and cell aggregates such as spheroids.

Computational approaches play a pivotal role in the strategic selection of cell strains for TERM biofabrication. Within this realm, cells are frequently sourced from the patient themselves, necessitating careful consideration regarding the types of cells to utilize. This involves making informed decisions about directing the differentiation processes of patient-derived induced Pluripotent Stem Cells (iPSC) towards specific cell types for regenerative applications. In this direction, computational methods proved capable of supporting the design of directed differentiation or transdifferentiation processes of cells. A computational framework that combines gene expression data with regulatory network information can predict the reprogramming factors necessary to induce cell transdifferentiation for TERM applications[130]. Furthermore, another computational framework is the base of a design tool for finding combinations of signals efficiently inducing cell conversions based on a stochastic gene regulatory network model embedding information on the transcriptional and epigenetic landscape of cells in [131].

Self-assembled 3D cell spheroids can replicate tissue functionality, working as building blocks for TERM products and supporting *in vitro* tissue maturation before fabrication by bioprinting or bioassembly, increasing the efficiency of overall fabrication processes of TERM products. Computational methods sustain the rational fabrication of spheroids to tune them to specific application needs. For example, computation combines with experimental activity to support the modulation of non-geometrical parameters in the design of the fabrication process of 3D articular chondrocytes spheroids. In contrast, CFD modeling underlies the optimization of geometrical features of the culture chip, modulation of cell-related features such as cell concentration, flow rate, and seeding time to model cell trapping follows in [132].

In conclusion, computational methods support biomaterials qualification for TERM biofabrication by sustaining the rational choice and combination of inorganic and organic constituents of biomaterials, taking into account biological requirements to optimize inorganic biomaterials, as well as the delicate nature of biological components in cell-laden constructs.

### 4.3. Fabrication

Fabrication of spatially-organized TERM constructs (Figure 12.C) employs raw cells and biomaterials and relies on various techniques. Among them, this review focuses on bioprinting and bioassembly. Bioprinting is *“the use of computer-aided transfer processes for patterning and assembling living and non-living materials with a prescribed two-or three-dimensional (2D or 3D) organization to produce bioengineered structures serving in regenerative medicine, pharmacokinetic and basic cell biology studies"* (adapted from [133]). Bioassembly is *“the fabrication of hierarchical constructs with a prescribed 2D or 3D organization through the automated assembly of pre-formed cell-containing fabrication units generated via cell-driven self-organization or through the preparation of hybrid cell-material building blocks, typically by applying enabling technologies, including microfabricated molds or microfluidics"* [1].

#### 4.3.1. Printing process

Bioprinting applies to both inorganic scaffolds and cell-embedded bioinks. The computational optimization of bio-printing processes potentially has various objectives [58]. Some endeavors aim to maximize printing fidelity: the similarity between the product models and the actual bio-printed product. Other approaches maximize biomimetic fidelity: the construct’s biological, mechanical, and rheological similarity to its *in vivo* counterpart. Other approaches tackle the challenge of the joint optimization of printing and biomimetic fidelity in bioprinting. For example, computational analysis underlies the tuning of flow parameters to optimize needle geometry for the biofabrication of high-resolution, fragile cell transfer of printed human iPSC with a low-cost 3D printer for precise cell placement and stem cell differentiation in [134].

Several computational DoE approaches target the optimization of significant process parameters in Fused Deposition Modeling (FDM), a rapid additive manufacturing prototyping technique [135]. DL methods proved able to optimize the bioprinting process by elucidating the complex relationships among the various printing parameters and predicting their optimal configurations for specific objectives [136, 137, 60]. For example, a multi-objective optimization approach improves printing accuracy and stability in [138]. In contrast, a ML approach underlies effective and accurate drop-on-demand printing control in [139]. Computational methods also sustain quality control in the printing process. For example, a CNN supports the data-driven detection of printing anomalies towards real-time process adjustment to ensure high printing quality in [140].

Computational optimization also targets the design of fabrication processes and systems. A fuzzy Technique for Order of Preference by Similarity to Ideal Solution (TOPSIS) Multi-criteria Decision Making (MCDM) process allows for identifying the best bioprinting process to optimize the biomechanical properties of the construct given a starting printing material in [141]. Multi-physics computational models support the design of an extruder system for the biofabrication of vascular networks, intending to maintain cell viability during the printing process [142].

#### 4.3.2. Cell seeding

Maximizing the number of cells uniformly occupying the available space within scaffolds facilitates tissue culture functionality and reduces tissue maturation time. Thus, cell seeding is a critical step in the biofabrication of TERM products. Computational methods combine with experimental activity to design effective cell seeding techniques. For instance, a combination of FEM to predict the scaffold permeability effects on cell seeding effectiveness and a fully factorial experimental design allowed to assess the relative importance of permeability, thickness, and coating on cell seeding efficiency and uniformity in [143]. As an additional example, the simulation of the cellular interactions with biomaterials, cells, and dynamic media in a scenario where cell seeding occurs under perfusion conditions helped to optimize the seeding time, and the number of cells seeded in the scaffold in [144].

#### 4.3.3. Spheroid bioassembly

Spheroid fusion, the aggregation of spheroids into larger constructs, is an approach for efficiently constructing larger tissues. With the increase in construct size, oxygen, and metabolites are no longer uniformly accessible to cells across the construct, hampering their viability and thus the quality of fabrication products. Computational methods support modeling cellular needs during spheroid-dependent fabrication, predicting their viability under different process parameters. For example, a hybrid discrete-continuous heuristic model, combining a cellular Potts-type approach with field equations, describes metabolic effects over cells during spheroid-dependent fabrication in [145].

In conclusion, computational methods sustain the multi-objective optimization of the fabrication stage of TERM biofabrication by navigating complex trade-offs between fabrication parameters, benefiting from the explicit modeling of the role of cells in ensuring final product quality. This advantage applies to both inorganic and organic components of fabricated products.

### 4.4. Maturation

Automated culture systems include different technologies ranging from microfluidic devices to bioreactors (Figure 12.D), which automatically administer signals to guide product maturation towards the desired structural and functional features. In biofabrication, the maturation stage is a critical stage where the biofabricated structures undergo cellular development, differentiation, and extracellular matrix formation to achieve functional characteristics akin to natural tissues [52]. The dynamical modulation of environmental stimuli administered to the construct affects product maturation and, ultimately, the emergence of the desired functional and structural features. Computational methods support the characterization and design of the maturation process, allowing prediction of how the administered stimuli make the desired features emerge [146]. For example, DL approaches to support the prediction of functional activations from environmental stimuli of different kinds [60]. ANNs predicted the induction of osteogenesis from sets of biomechanical loading parameters values [147]. Computational approaches also target the overall biofabrication process design by predicting the final product features and the respective product quality based on the set of administered stimuli and their organization in time and space.

For example, modeling the identity and quality of stimuli, including their complex behavior in the physical environment, combines with experimentation to link biological responses to mechanical stimulation, improving the understanding of the cause-effect relationship of mechanical loading in [148].

As a notable computational approach, SIMulation using Metropolis Monte Carlo method (SIMMMC) supports the 3D modeling of specific biological systems, including living cells and biomaterials, and simulates construct evolution from the structural and functional perspective, based on the Metropolis Monte Carlo (MMC) modeling of living cells, biomaterials, and cellular medium [149].

Maturation often relies on automated culture systems such as bioreactors of varying sizes and designs. Computational methods support the optimization of bioreactor design. For example, CFD modeling allows the optimization of the flow profile when designing the perfusion chamber of a perfusion bioreactor system to engineer human cartilage grafts in [150].

Construct maturation’s success in bioreactors relies heavily on cell viability and proliferation. Computational models of the micro- and the macro-scale combine with CFD to simulate cell colonization processes in a perfusion bioreactor and perform model-based optimization of the perfusion flow rate to maximize cell colonization in [151]. A model of tissue growth inside 3D scaffolds in a perfusion bioreactor combines with a multi-objective optimization method to find the most cost-effective medium refreshment strategy for maximizing tissue growth while minimizing experimental costs in [152]. Discrete simulation of intra- and extracellular processes combines with evolutionary computation to perform the DSE of process designs to generate optimal biofabrication protocols to maximize size and control geometry of human epithelial monolayers in silico in [153]. Continuous simulation of tissue dynamics based on vertex models of cells (leveraging on the PalaCell2D simulation framework [154]) combine with a deep Reinforcement Learning (RL) approach to generate optimal protocols for epithelial sheets culture for maximizing the total number of cells produced and optimizing the spatial organization within the cell aggregate in [155].

In conclusion, computational methods support the maturation stage of TERM biofabrication by modeling the dynamic processes taking place within the developing construct and predicting the effect of the administered stimuli over them to explore several process outcomes and support the choice of optimal process designs for specific target products.

## 5. Open challenges and future trends

This review aimed to present computational methods for modeling, designing, and optimizing TERM biofabrication processes, with a focus on encompassing biological complexity through diverse modeling approaches from holistic to hypothesis-driven. The predominance of TERM bone or cartilage engineering in publications aligns with science mapping results (Section 2), while the infrequent application in more complex tissue or organ biofabrication reflects current limitations in mimicking biological complexity (section 1). Despite these limitations, computational tools show promise in advancing the biofabrication of complex TERM products, thus expanding the range of regenerative applications. However, current computational models often simplify the intricate relationship between organic and inorganic components in biofabrication or focus narrowly on single process aspects, limiting their impact on process design and product quality. To fully realize the potential of computational optimization in TERM biofabrication, a truly interdisciplinary approach is necessary, incorporating diverse scientific and technological elements and addressing the underlying biological complexity for holistic process modeling, design, and optimization, while developing strategies to improve their computational feasibility. In particular, future computational methods in TERM biofabrication should address the following key challenges.

### Interdisciplinary enabling

Computational methods must become facilitators of interdisciplinary collaboration, catering to the varied scientific backgrounds of researchers in TERM biofabrication [1]. They should ensure usability and accessibility, promoting findability and direct reusability of solutions [156] to accelerate collaborative scientific advancement. The sharing of novel solutions should leverage on scientific reproducibility tools like Docker or Singularity container platforms [157]. In general, novel methods shall truly enable interdisciplinary collaboration, to mediate the synthesis of diverse perspectives into models, and providing modular and reusable solutions for model-based optimization.

### Intrinsic flexibility

Computational methods must adapt to new biofabrication processes and targets, especially in optimizing clinically relevant products. They should support personalized medicine principles, adapting to specific patient needs [158, 159, 160] and the complex normative scenario for clinical translation [161, 162]. Future approaches shall reach extensive flexibility over of patient- and regulation-dependent constraint specifications to seamlessly adapt to different clinical applications.

### Multi-component modeling

computational methods must rely on accurate models of the several inorganic and biological components in biofabrication [2, 6], accurately predicting cellular behavior during fabrication and maturation, and sustaining the optimization of the process design accordingly [9]. Multi-component models can rely on diverse approaches on the broad spectrum between holistic, comprehensive modeling and hypothesis-driven modeling [49]. Future approaches shall find the right trade-off between these two, or sustain hybrid modeling, consistently combining model components based on different modeling strategies [50].

### Multi-stage modeling

computational methods must comprise the multiple process stages composing a TERM bio-fabrication process [163, 164], drawing meaningful relations between them to support multi-stage modeling and optimization. In this perspective, computational models shall either model multiple stages together, or support the usage of hybrid models [50] that cover multiple stages by composition of multiple single-stage models.

### Abstraction tuning

Computational methods must develop to sustain tunable abstraction in TERM biofabrication models. The abstraction of mechanistic models into behavioral models, while preserving good model prediction accuracy and explainability, is an established strategy to enhance computational feasibility of model-based DSE in other domains [165]. Future research shall implement the same principle to sustain tunable abstraction in model-based DSE of TERM biofabrication processes.

### Computational scale-up

Computational methods must enhance the feasibility of model-based DSE for TERM biofabrication, especially when dealing with large design spaces. Heuristic and meta-heuristic optimization strategies in model-based DSE can efficiently explore huge design spaces [43]. Novel computational methods shall implement and innovate such techniques towards a general increase in the exploration of large TERM biofabrication process design spaces.

These advances will make computational approaches the key for unlocking the potential of TERM biofabrication for faster clinical translation, targeting more complex applications and reaching improved clinical outcomes.

### Competing interest statement

The authors declare that they have no known competing interests that could have appeared to influence the work reported in this paper.

## Funding acknowledgement

This study was carried out within the project "SAISEI - Multi-Scale Protocols Generation for Intelligent Biofabrication" funded by the Ministero dell’Università e della Ricerca (Italian Ministry for Universities and Research) – within the Progetti di Rilevante Interesse Nazionale (PRIN) 2022 program (D.D.104-02/02/2022).

## CRediT authorship contribution statement

**Roberta Bardini:** Conceptualization; Data curation; Investigation; Methodology; Visualization; Writing - original draft; Writing - review and editing.. **Stefano Di Carlo:** Conceptualization; Funding acquisition; Project administration; Supervision; Writing - review and editing..

## Abbreviations

2D: two-dimensional
3D: three-dimensional
4D: four-dimensional
3D-CNN: Three Dimensional Convolutional Neural Network
AI: Artificial Intelligence
ANN: Artificial Neural Networks
BESO: Bidirectional Evolutionary Structural Optimization
BHV: Bioprosthetic heart valve
CAD: Computer-Aided Design
CAE: Computer-Aided Engineering
CAM: Computer-Aided Manufacturing
CFD: Computational Fluid Dynamics
CT: Computed Tomography
CNN: Convolutional Neural Network
FMEA: Failure Modes and Effects Analysis
DICOM: Digital Imaging and Communications in Medicine
DL: Deep learning
DoE: Design-of-Experiments
DSE: Design Space Exploration
DT: Digital Twin
ECM: Extra-Cellular Matrix
FDM: Fused Deposition Modeling
FE: Finite Elements
FEM: Finite Elements Methods
FSI: Fluid-Structure Interaction
IDT: Intelligent Digital Twin
iPSC: induced Pluripotent Stem Cells
MADM: Multi-attribute Decision Making
MCDM: Multi-criteria Decision Making
MMC: Metropolis Monte Carlo
MODM: Multi-objective Decision Making
ML: Machine Learning
MRI: Magnetic Resonance Imaging
OFAT: One Factor at A Time
QFD: Quality Function Deployment
QbD: Quality by Design
RL: Reinforcement Learning
RP: Rapid Prototyping
RPA: Robotic Process Automation
SIMP: Solid Isotropic Material with Penalization
STL: Standard Tassellation Language
TO: Topology Optimization
TOPSIS: Technique for Order of Preference by Similarity to Ideal Solution
TPMS: Triply periodic minimal surfaces
SIMMMC: SIMulation using Metropolis Monte Carlo method
TERM: Tissue Engineering and Regenerative Medicine
TP: Total Publications

---

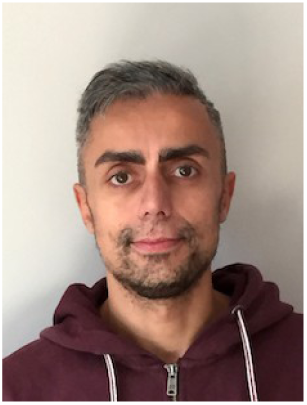
Stefano Di Carlo is a full professor in the Control and Computer Engineering department at Politecnico di Torino (Italy) since 2021. He holds a Ph.D. (2003) and an M.S. equivalent (1999) in Computer Engineering and Information Technology from the Politecnico di Torino in Italy. Di Carlo’s research contributions include biological network analysis and simulation, machine learning, image processing, and evolutionary algorithms, as well as Reliability, Memory Testing, and BIST.

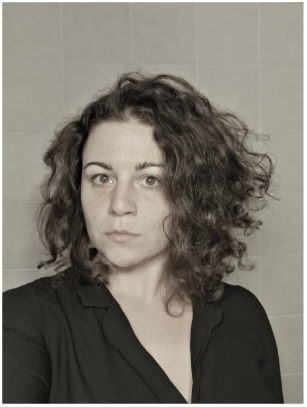
Roberta Bardini is a post-doc researcher at the Department of Control and Computer Engineering of Politecnico di Torino. She holds a Ph.D. (2019) in Control and Computer Engineering from the Politecnico di Torino in Italy, and a M.Sc. in Molecular Biotechnology (2014) from Università di Torino. Her research focuses on computational biology and bioinformatic approaches for analyzing and modeling complex biological systems, and on the computational design and optimization of biomanufacturing processes.

## Footnotes

1 https://pubmed.ncbi.nlm.nih.gov/

## Notes

### Competing Interest Statement

The authors have declared no competing interest.

### Summary of Updates

his version of the manuscript has been revised to update the sections organization and paragraph structure after a major revision.

